# Comparing Automated Subcortical Volume Estimation Methods; Amygdala Volumes Estimated by FSL and FreeSurfer Have Poor Consistency

**DOI:** 10.1101/2024.03.07.583900

**Authors:** Patrick Sadil, Martin A. Lindquist

**Affiliations:** Johns Hopkins Bloomberg School of Public Health

**Keywords:** MRI, Amygdala

## Abstract

Subcortical volumes are a promising source of biomarkers and features in biosignatures, and automated methods facilitate extracting them in large, phenotypically rich datasets. However, while extensive research has verified that the automated methods produce volumes that are similar to those generated by expert annotation, the consistency of methods with each other is understudied. Using data from the UK Biobank, we compare the estimates of subcortical volumes produced by two popular software suites: FSL and FreeSurfer. Although most subcortical volumes exhibit good to excellent consistency across the methods, the tools produce diverging estimates of amygdalar volume. Through simulation, we show that this poor consistency can lead to conflicting results, where one but not the other tool suggests statistical significance, or where both tools suggest a significant relationship but in opposite directions. Considering these issues, we discuss several ways in which care should be taken when reporting on relationships involving amygdalar volume.

## 1. Introduction

Regional volumes of subcortex have been proposed as biomarkers for several psychopathologies. For example, the volume of the amygdala has been suggested as a biomarker for Alzheimer’s, depression symptom severity in young adults, bipolar disorder in youth, migraine frequency, chronic pain, and others (Daftary et al., 2019; Khatri and Kwon, 2022; Pfeifer et al., 2008; Liu et al., 2017; Vachon-Presseau et al., 2016; Rogers et al., 2009; Ruocco et al., 2012; Szeszko et al., 2004). As a biomarker, subcortical volumes are advantageous for being interpretable (given the rich literature linking these structures to many functions), explainable (hypotrophy and hypertrophy are both easily described to healthcare providers and patients), and readily available.

The latter point comes from the fact that it is possible to estimate the regional volumes from any structural image with several automated algorithms.

For estimating regional subcortical volumes, two automated techniques are popular: FMRIB’s Integrated Registration and Segmentation Tool (FIRST) from the FMRIB Software Library (FSL) and FreeSurfer’s Automated Segmentation (ASEG) (Patenaude, 2007; Patenaude et al., 2011; Fischl, 2012). Both techniques exhibit high consistency with the gold-standard of manual segmentation in healthy adults and some clinical populations (Hsu et al., 2002; Tae et al., 2008; Morey et al., 2009; Pardoe et al., 2009; Dewey et al., 2010; Lehmann et al., 2010; Doring et al., 2011; Nugent et al., 2013; Wenger et al., 2014), although there is variability across regions and between methods. For segmenting the hippocampus, FreeSurfer has been reported as having higher intraclass correlations than FSL (Doring et al., 2011), and neither method appears to have worse reliability (across repeated scans) than manual segmentation (Mulder et al., 2014). For the putamen, FSL has a higher Dice coefficient with manual segmentation, and the methods perform similarly on the caudate (Perlaki et al., 2017). For the amygdala, which method performs better depends on the metric (Morey et al., 2009). However, these comparisons may not generalize to other populations (e.g., pediatric, elderly), given that performance of the automated techniques is not consistently high across populations (Schoemaker et al., 2016; Kim et al., 2012; Sánchez-Benavides et al., 2010; Zhou et al., 2021). Both tools may remain popular because their performance against a gold-standard depends on the research context, with different situations favoring different tools.

Given that both tools are often reasonable choices for a given study, and that the literature contains many reports that are based on one but not the other, we sought to understand how the tools compare with each other. To our knowledge, the two tools have only been compared to each other by Perlaki et al. (2017). In their research, the tools were consistent with each other (exhibiting intraclass correlations that ranged from around 0.7 - 0.9), but the analyses included only the putamen and caudate, and the sample size was relatively small (N=30). That is, it remains unclear how often the two tools provide the same results and which factors affect these discrepancies.

We extend the results of Perlaki et al. (2017) to the remaining subcortical structures using a much larger population (tens of thousands). Our investigation should be considered in the context of research that uses estimates of volume as a biomarker by, for example, correlating it with a health-related outcome. Our primary concern is whether the results of such a study are expected to depend on the method used for automated segmentation.

## 2. Methods and Results

First, we looked at the consistency of subcortical volumes between FSL and FreeSurfer using data from the UK Biobank (Alfaro-Almagro et al., 2018). The data were downloaded Jan 2024 and contained 45743 participants with usable anatomical data (Category 190: FreeSurfer ASEG; Category 1102: FSL FIRST). Agreement and consistency of the tools were measured using the single-measurement intraclass correlation coefficient. For details on intraclass correlation calculations, see Appendix A.1. Estimates and associated uncertainty are displayed using subscripts, as recommended by Louis and Zeger (2008). For example, an estimate of 0.22 with a 95% confidence interval spanning [0.21, 0.23] will be rendered as _0.21_0.22_0.23_.

Across all structures, estimated agreement was lower than estimated consistency (Table A1), reflecting differences in the average volumes estimated by the two methods (Appendix A.2). However, a constant shift across participants would not affect many analyses targeted by our primary concern, analyses related to estimated correlations between regional volumes and some other measure. For that reason, we focus not on agreement but instead on consistency.

Consistency varies by region (Figure 1, Table A1). To interpret the intraclass correlations, consider the categories provided by Cicchetti and Sparrow (1981): <0.4: poor, [0.4, 0.6): fair, [0.6, 0.75): good, [0.75, 1): excellent. Using those categories, the methods exhibit “good” to “excellent” consistency for most regions. However, the consistency of volumes for the amygdala is markedly worse than the others, being around only _0.24_0.24_0.25_ and _0.21_0.22_0.23_ for the left and right hemispheres (ranges of uncertainty span 95% confidence intervals). Compare those values to the values for the hippocampus (Figure 1), which has good consistency (left: _0.69_0.69_0.70_, right: _0.70_0.70_0.71_). For both structures (that is, the amygdala and hippocampus), consistency across hemispheres as reported by FreeSurfer is the highest numerically (Figure 1).

**Figure 1:**
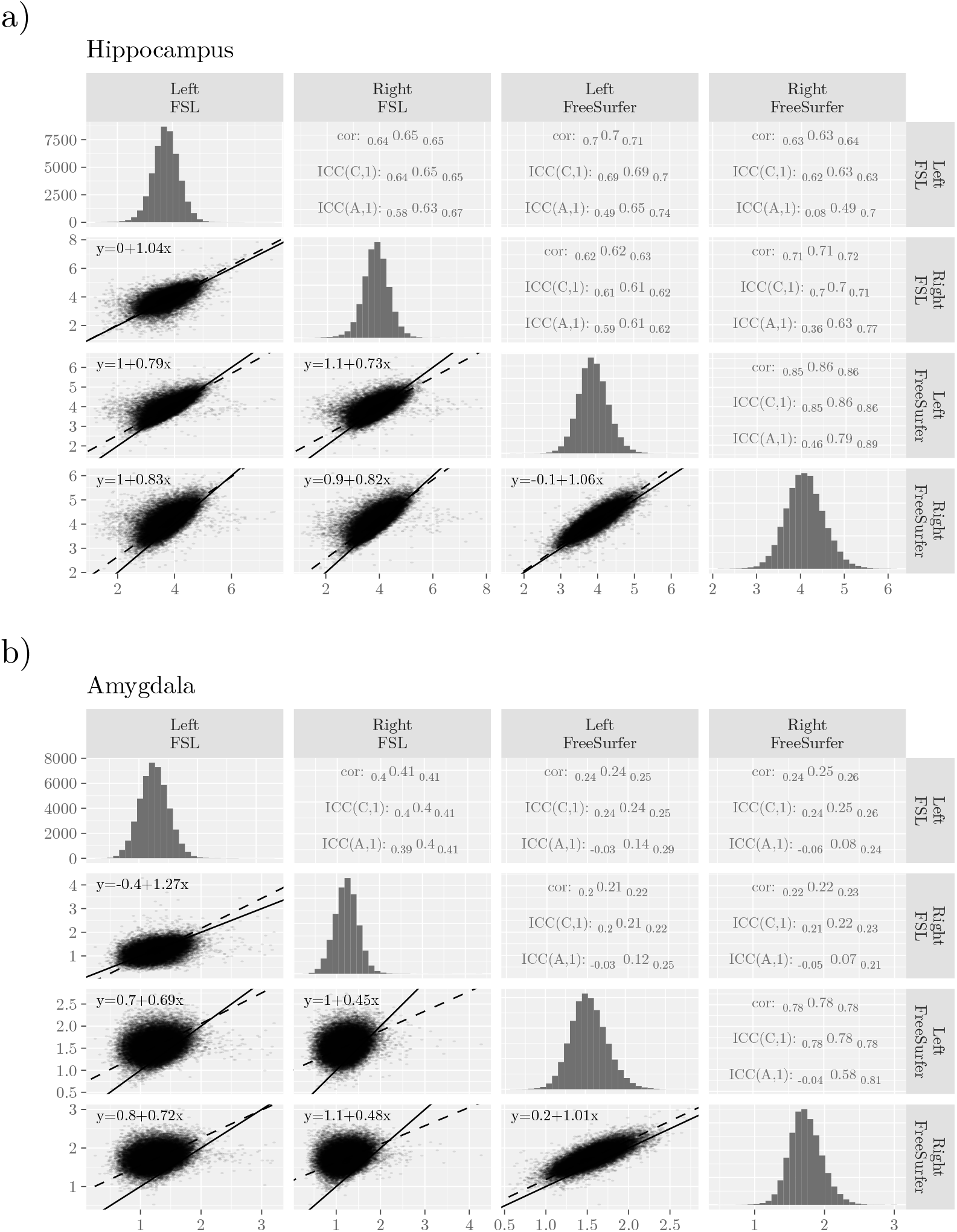
Comparisons of Subcortical Volumes Estimated by FSL and FreeSurfer for two example regions Hippocampus and b) Amygdala. For remaining subcortical regions, see Table A1. In the lower triangular panels, the line of equivalence is marked with a solid line, and the dashed lines show the result of orthogonal regression. In the upper panel, the uncertainty estimates span 95% confidence intervals. The histograms along the diagonal display volumes in the full sample.

**Figure 2:**
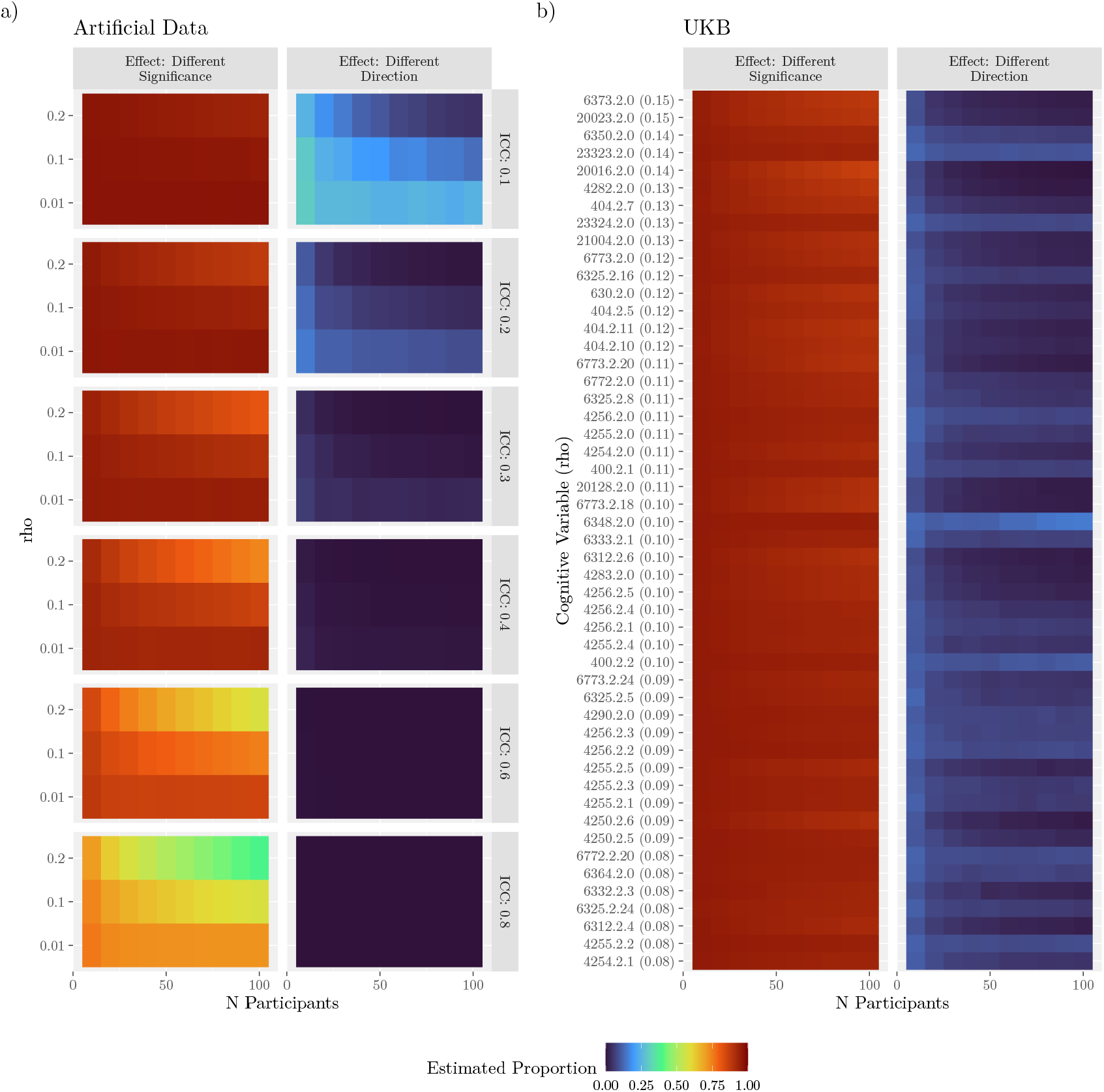
Effects of Low Measurement Consistency. In the subfigures, the left column or panel shows the proportion of simulated experiments where the methods produce correlations on opposite sides of an 𝛼 = 0.05 significance threshold. The right column or panel shows the proportion of simulated experiments where both measures are significantly correlated with an outcome but in opposite directions. Proportions are shown with color (Swihart et al., 2010). a) Simulations with Artificial Data. Rows (rho) indicate pre-specified product-moment correlation with the outcome variable. ICC: pre-specified intraclass correlation between measures of volume. b) Simulations with UKB. Each row corresponds to a non-image derived phenotype from the UKB. The value in parentheses is the absolute correlation of the variable with the average volumes of the left amygdala (average across methods). For a display without color that includes estimates of uncertainty, see Figure A3.

Although our focus is on the consistency between tools, we note that their lack of agreement may lead researchers to make conclusions about whether one hemisphere tends to have a larger amygdala than the other. There is an interaction between method and hemisphere (FSL - FreeSurfer: 0.22, 𝑝 < 0.001), with FreeSurfer reporting that the right amygdala is larger than the left (left - right: -0.19, 𝑝 < 0.001) and FSL reporting that the left is larger than the right (left - right: 0.04, 𝑝 < 0.001). As the effect side is almost zero in the latter, practically the interaction is driven by the right side.

The amygdala has been described as particularly challenging to segment; one small study (N=23) reports a consistency of 0.6 for volumes estimated by FSL across repeated scans of the same individual (Morey et al., 2010). Moreover, automated segmentation algorithms can be affected by experimental factors like site, scanner, participant positioning, and software version (Hedges et al., 2022; Du et al., 2021; McGuire et al., 2017; Yang et al., 2016; Liu et al., 2020; Mulder et al., 2014; Morey et al., 2010; Perlaki et al., 2017), and differential sensitivity to such factors could impact an intraclass correlation. In the UKB, several potentially confounding factors were correlated with estimates of amygdalar volume (Figure A2). However, regressing these factors from the estimates of volume did not improve the consistency between the methods (Appendix A.4).

**Figure A2:**
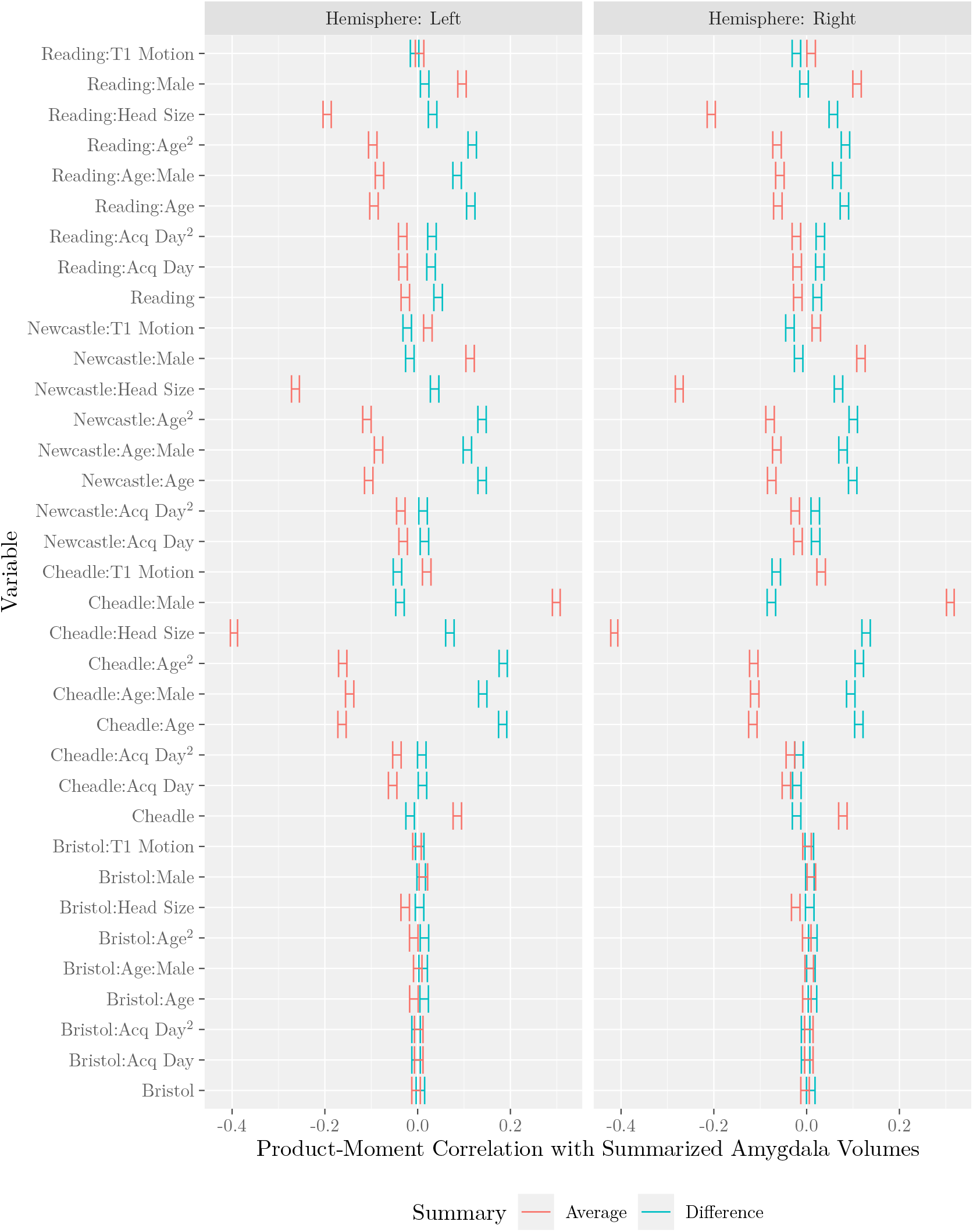
Correlation Between Potential Confounders and Amygdala Volume Measurements. Summary describes how the the volumes were combined across methods. Summaries were either an average or a difference (FSL-FreeSurfer). Note that the variable “Head Size” corresponds to a scaling factor, and that larger values imply smaller brains.

With poor consistency between measurements of the amygdala, there is concern that reported relationships involving amygdala volumes may depend on which method is used for estimating the volume, a choice that may be considered arbitrary or lab-specific. At least two kinds of issues could arise. First, lower consistency could make it more likely that one but not both methods leads to significant correlations. Second, lower consistency could make it more likely that the two methods produce significant correlations that go in opposite directions.

To investigate how often these two issues could occur, we first simulated experiments with artificial data. In each simulation, datasets with two noisy estimates of volume and a third, outcome, variable were generated such that the two volume estimates had a prespecified intraclass correlation with each other (in expectation), and the true volume had a given product-moment correlation with the outcome variable. The estimated volumes were then tested for a product-moment correlation with the outcome, and the process was repeated for several intraclass correlations and sample sizes. In all simulations, the true product-moment correlation was set to a value that is either typical for neuroimaging research (0.1, Marek et al., 2022), small but non-zero (0.01), or large (0.2). For additional details on the simulation methods, see Appendix A.5.

With lower consistency, the estimated product-moment correlations were more often on opposite sides of a significance threshold (Figure 2a, left column). For example, with consistency near the value that was estimated for volumes of the amygdala in the UKB (0.2), a typical effect size (0.1), and experiments with 50 participants, significance differed in around _94.9_95.0_95.1_ percent of simulations in which one test was significant (for simulations, estimates indicate medians and ranges of uncertainty span 95% equal-tailed intervals, see Appendix A.5). With a sample size of 100, that percentage was only minimally different (to _93.7_93.9%_94.0_). For a discussion on this insensitivity to sample size, see Section Appendix A.6.

Lower consistency also coincided with a higher proportion of experiments in which the two methods correlate in opposite directions (Figure 2a, right column). Considering the previous example (an intraclass correlation of 0.2, an effect size of 0.1, and 50 participants per experiment), around _5.13_5.71_6.33_ percent of experiments in which both product-moment correlations were significant resulted in the correlations having opposite signs. As the intraclass correlation increased, the rates of this effect decreased rapidly.

To assess how often these two issues could occur in practice, we repeated the above analyses but with the UKB. From the UKB, we extracted the “Cognitive” variables using the FMRIB UKBiobank Normalisation, Parsing And Cleaning Kit (McCarthy, 2023), restricting analyses to variables with instance 2 and that exhibited one of the largest (in absolute magnitude) 50 rank correlations (between the cognitive variables and the average of the two estimates of the volume of the left amygdala), and further to only those participants that had values for all 50 of those variables (23486 participants). Each simulation resulted in a rank correlation between the two volume estimates for the left amygdala and each of the 50 variables (results were comparable with the right amygdala). For additional details, see Appendix A.5.

Considering differing significance levels, the rates across experiments using the UKB resembled the rates from experiments with artificial data. In simulated experiments with 50 UKB participants in which at least one correlation was significant, one correlation was not significant in between _90.7_90.8_91.0_ and _95.1_95.2_95.3_ percent of simulations (the two estimates cover the 50 cognitive variables; Figure 2b, left panel). Proportions were generally lower for variables that were more strongly correlated with the measure of volume (for illustration, compare the variables that are higher versus lower in the left panel of Figure 2b).

Considering significant correlations with differing signs, the rates across experiments with the UKB bracketed the rates with artificial data. In experiments simulated with 50 UKB participants, the rates ranged from _1.28_1.45_1.64_ to _7.64_8.32_9.03_. For most variables, increasing the number of participants decreased the proportion of experiments in which the two correlations exhibited opposite signs, but for some the proportion increased. Across the 50 variables, 9 had a higher proportion at N=100 than N=50, 2 of which had non-overlapping 95% equal-tailed intervals (6348: duration to complete numeric path; 400: time to complete round of pairs matching game; see also Figure A3).

**Figure A3:**
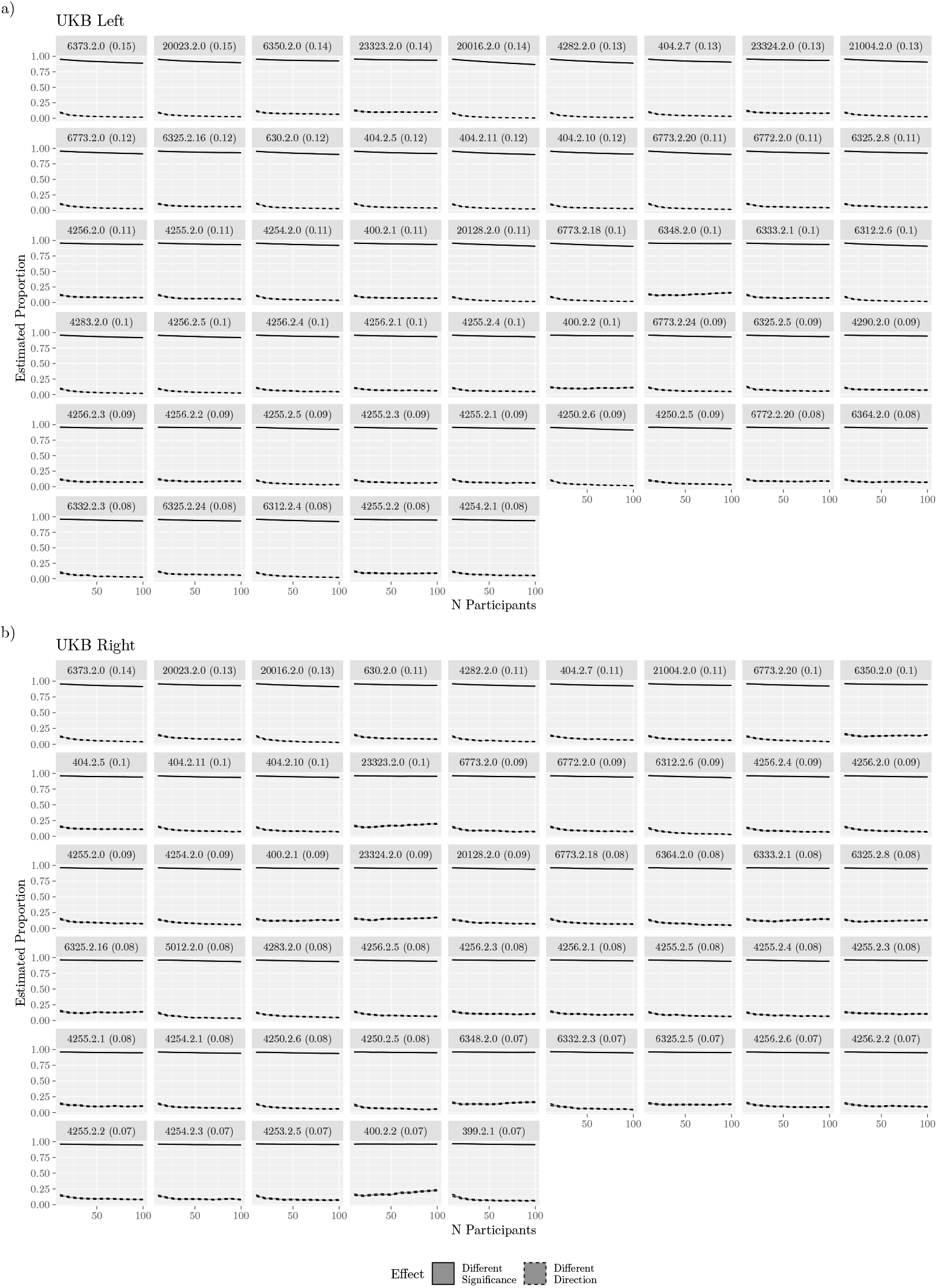
Effects of Low Measurement Consistency on the UKB in Left and Right Hemispheres. Rib- bons span 95% equal-tailed interval estimated from simulated experiments. See also Figure 2, where the medians are represented with color.

## 3. Discussion

We examined the consistency of subcortical volumes within the UK Biobank (Alfaro-Almagro et al., 2018), observing that two common methods of estimating the volume of the amygdala, one from FSL and one from FreeSurfer, have poor consistency with each other.

We speculate that the lack of consistency for the amygdala compared to other subcortical structures is due in part to its relatively small size. For larger structures, discrepancies in a few voxels would have a smaller impact on consistency as compared to the impact of the same discrepancy on a smaller structure. When considering the difference in amygdala volume estimates between tools as a proportion of their average (Figure A1 b), a trend is revealed where smaller structures tend to have more variable differences than larger ones. The amygdala is the smallest structure analyzed and has the largest variation. Even within other structures, the variability of the proportional difference increases as the average size decreases. Moreover, there is a clear correlation between head size and difference in amygdala volume (Figure A2), whereby smaller heads are associated with larger differences. Hence, smaller structures may be more difficult to segment consistently due to the proportional impact of discrepancies in the classification of a few voxels.

The main concern in this report is that consistency this poor can lead to conflicting results. Two kinds of conflict were explored: the methods producing correlations that are on opposite sides of significance thresholds, and the methods producing volumes with significant correlations that have differing signs. The prevalence of these occurrences was estimated with artificial data and data from the UKB. Based on the observed rates, we make the following recommendations.

### 3.1. Recommendations

#### 3.1.1. When testing for new biomarkers, report relationships with multiple automated methods (e.g., both FSL and FreeSurfer)

Researchers may have idiosyncratic reasons for selecting a method, particularly when the choice is viewed as arbitrary. If the choice between methods is arbitrary, then reporting the outcome across selections clarifies the fragility or robustness of a result (for a general discussion, see Steegen et al., 2016). The choice may not always be arbitrary, as there are metrics along which and study populations for which one method may perform better (Zhou et al., 2021; Morey et al., 2009; Huizinga et al., 2021). Note that one effect of low consistency could be a downward bias on the magnitude of estimated correlations (Section Appendix A.7), and so there may be advantages to not only reporting but also combining the estimates across methods when predicting health-related outcomes (e.g., by averaging, or including both as independent variables in predictive models). Reporting estimates from multiple reasonable tools will help move conclusions beyond “there exists a correlation with the volume of the amygdala as estimated by method M (version x)” to simply “there exists a correlation with the volume of the amygdala”.

#### 3.1.2. When reviewing or conducting meta-analyses of relationships with amygdala volume, consider the method that was used to estimate volume

As mentioned in the Introduction, the amygdala has received substantial attention due to being predictive of an array of health-related outcomes. As presented in this report, the volume that is estimated by one automated method may only weakly correspond to the volume estimated by another, and so it may be misleading to conduct a meta-analysis without accounting for the algorithms used by the individual studies. In the UKB, there are differences in the strength of the correlations between the measures of volume and the cognitive variables; in nearly all of the variables considered in this report, the magnitude of the correlations involving FreeSurfer’s method were numerically larger (Figure A5). Larger correlations do not imply higher veracity, but they further indicate that the methods track aspects of amygdalar anatomy differently.

#### 3.1.3. When replicating or extending research on a relationship that involves the volume of the amygdala, use the method reported in the original publications

This recommendation follows standard practice for a replication study. We highlight it here in consideration of both extension studies that aim to apply a biomarker or biosignature that includes amygdala volume (such as when testing a putative relationship in a new population), and also in consideration of the ongoing evolution of methods for automatically estimating subcortical volumes. Although FSL and FreeSurfer are two of the most popular methods, others exist (e.g., Akhondi-Asl and Warfield, 2013), including newer techniques based on deep-learning approaches (e.g., Billot et al., 2023; for review, see Singh and Singh, 2021). Newer methods may have better correspondence with manual segmentation, warranting their use in replication or extension studies. But as this report shows, two methods can perform well while exhibiting poor consistency with each other. So when building on prior findings, it remains important to use the methods of those prior findings, even when they are superseded.

## Code Availability

Code to reproduce analyses is available on GitHub: https://github.com/psadil/auto-volume-comparisons.

## Author Contributions

**Patrick Sadil**: Conceptualization, Methodology, Software, Validation, Formal Analysis, Investigation, Resources, Data Curation, Writing - Original Draft, Writing - Review & Editing, Visualization. **Martin A. Lindquist**: Conceptualization, Methodology, Validation, Formal Analysis, Resources, Writing - Original Draft, Writing - Review & Editing, Supervision, Project Administration, Funding Acquisition.

## Ethics Statement

Informed consent was obtained from all UK Biobank participants. Ethical procedures are controlled by a dedicated Ethics Advisory Committee (http://www.ukbiobank.ac.uk/ethics).

## Acknowledgements

This work was supported by grant R01MH129397 from the National Institute of Mental Health. This research has been conducted using data from UK Biobank, a major biomedical database (Project ID: 33278). We are grateful to UK Biobank and the UK Biobank participants for making the resource data possible, and to the data processing team at Oxford University for sharing the processed data. The UK Biobank imaging project is funded by the Medical Research Council and the Wellcome Trust.

## Appendix A. Supplementary Materials

### Appendix A.1. Intraclass Correlation

The intraclass correlation was based on a two-way mixed linear model (McGraw and Wong, 1996). In the model, the volume 𝑥 for the region of participant 𝑖 as measured by method 𝑘 was treated as equal to the sum of an intercept, 𝜈, the “true” volume, 𝜆_𝑖_, a method bias, 𝑐_𝑘_, and an error term, 𝜖_𝑖𝑘_

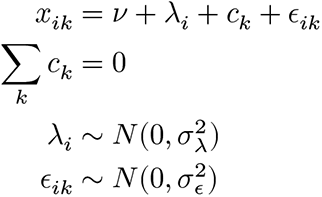

Note that the 𝑐_𝑘_ terms are assumed fixed, with variance given by 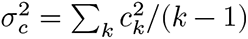

An important assumption of this model is that the two methods are expected to have the same mean-squared error. The model does not include features that would allow for the errors in the two methods to be correlated (e.g., participants are not distinguished by characteristics that coincide with the methods performing better or worse).

The consistency version of the intraclass correlation, 𝐼𝐶𝐶(𝐶, 1), and the absolute agreement, 𝐼𝐶𝐶(𝐴, 1) were given as fractions of the variance components

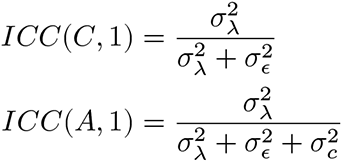

Variance components and associated confidence intervals estimated using the `R` package `irr` (R Core Team, 2023; Gamer et al., 2019), which uses the mean square approach described by McGraw and Wong (1996).

### Appendix A.2. Differences in Average Volumes

Although we primarily focused on the consistency of the methods, we also observed shifts in the average volumes reported by the two methods (Figure A1). Differences in averages across methods have been reported previously (Gomez-Ramirez et al., 2022; Perlaki et al., 2017; Dewey et al., 2010; Huizinga et al., 2021). FSL tends to report volumes that are larger than those from FreeSurfer for the Accumbens, Amygdala, Hippocampus, and Pallidum, whereas for the Caudate, Putamen, and Thalamus FSL tends to report values that are smaller than those from FreeSurfer (all 𝑝 < 0.0001 for two-sided t-test). Across structures, the variability in the difference appears to covary with the average of the two volume estimates, increasing as the average volume estimate decreases (Figure A1 b).

**Figure A1:**
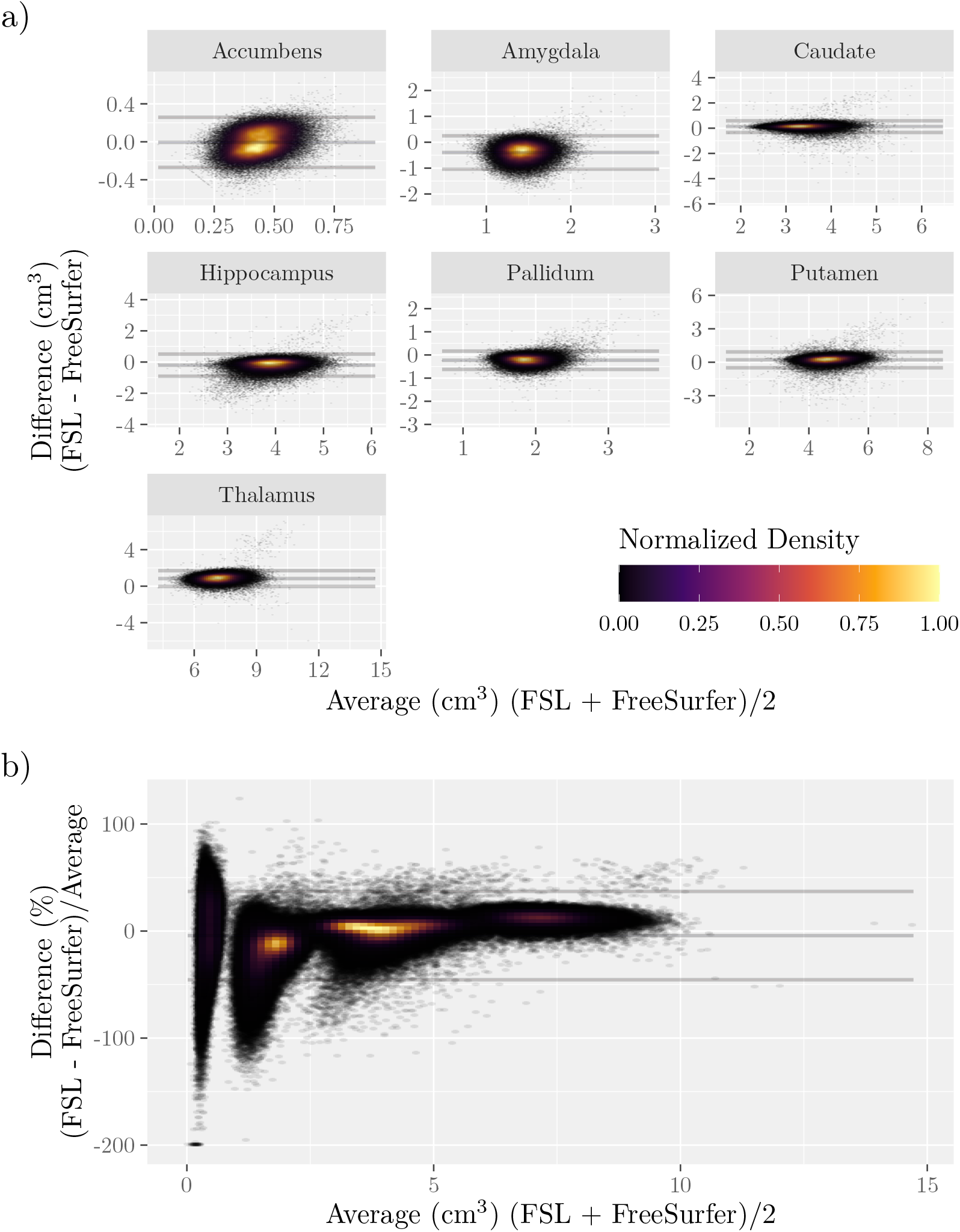
Bland-Altman Plots for Subcortical Volume. Horizontal lines show average difference and limits of agreement (1.96 standard deviations), with ribbons marking 95% confidence intervals. Panels correspond to subcortical structures. Left and right structures are plotted together. Shifts in the average estimate correspond to the central ribbon excluding zero. The color overlay indicates the degree of overplotting. a) Raw Differences. Note that axes are across panels independently. b) Differences by Percent Average.

**Table A1:** Intraclass Correlation for Subcortical Structures. Subscripts indicate 95% confidence intervals.

| Structure | Hemisphere | ICC(C,1) | ICC(A,1) |
| --- | --- | --- | --- |
| Accumbens | Left | 0.57<sup>0.58</sup><sub>0.58</sub> | 0.05<sup>0.44</sup><sub>0.65</sub> |
| Accumbens | Right | 0.56<sup>0.57</sup><sub>0.57</sub> | -0.03<sup>0.38</sup><sub>0.63</sub> |
| Amygdala | Left | 0.24<sup>0.24</sup><sub>0.25</sub> | -0.03<sup>0.14</sup><sub>0.29</sub> |
| Amygdala | Right | 0.21<sup>0.22</sup><sub>0.23</sub> | -0.05<sup>0.07</sup><sub>0.21</sub> |
| Caudate | Left | 0.85<sup>0.85</sup><sub>0.86</sub> | 0.75<sup>0.83</sup><sub>0.88</sub> |
| Caudate | Right | 0.86<sup>0.86</sup><sub>0.87</sub> | 0.62<sup>0.82</sup><sub>0.90</sub> |
| Hippocampus | Left | 0.69<sup>0.69</sup><sub>0.70</sub> | 0.49<sup>0.65</sup><sub>0.74</sub> |
| Hippocampus | Right | 0.70<sup>0.70</sup><sub>0.71</sub> | 0.36<sup>0.63</sup><sub>0.77</sub> |
| Pallidum | Left | 0.68<sup>0.68</sup><sub>0.69</sub> | -0.09<sup>0.41</sup><sub>0.71</sub> |
| Pallidum | Right | 0.66<sup>0.67</sup><sub>0.67</sub> | 0.04<sup>0.51</sup><sub>0.73</sub> |
| Putamen | Left | 0.79<sup>0.79</sup><sub>0.80</sub> | 0.54<sup>0.74</sup><sub>0.84</sub> |
| Putamen | Right | 0.82<sup>0.83</sup><sub>0.83</sub> | 0.56<sup>0.77</sup><sub>0.87</sub> |
| Thalamus | Left | 0.82<sup>0.82</sup><sub>0.83</sub> | -0.08<sup>0.49</sup><sub>0.79</sub> |
| Thalamus | Right | 0.83<sup>0.83</sup><sub>0.83</sub> | -0.09<sup>0.51</sup><sub>0.81</sub> |

### Appendix A.3. Intraclass Correlations Across All Subcortical Regions

#### Appendix A.4. Intraclass Correlations After Residualizing on Potential Confounds

One possible source of low intraclass correlations could be systematic inaccuracies with certain kinds of participants. For example, one of the two methods could tend to under- estimate volumes when given brains that have experienced severe atrophy, which would decrease the agreement of the methods. To assess this, correlations between the amygdala volumes and the “simple” confounds within the UKB were calculated (Alfaro-Almagro et al., 2021).

Several of the correlations appeared to be non-zero (Figure A2). However, regressing these confounds from the estimated amygdala volumes did not improve the intraclass correlations (Table A2).

**Table A2:** Intraclass Correlation After Deconfounding. Prior to calculating consistency, volumes were deconfounded by a version of the “simple” parameter set described by Alfaro-Almagro et al. (2021). For the list of variables, see Figure A2. Subscripts indicate 95% confidence intervals.

| Structure | Hemisphere | ICC(C,1) | ICC(A,1) |
| --- | --- | --- | --- |
| Accumbens | Left | 0.42<sup>0.43</sup><sub>0.44</sub> | 0.01<sup>0.30</sup><sub>0.51</sub> |
| Accumbens | Right | 0.44<sup>0.45</sup><sub>0.45</sub> | -0.05<sup>0.27</sup><sub>0.51</sub> |
| Amygdala | Left | 0.07<sup>0.08</sup><sub>0.09</sub> | -0.01<sup>0.04</sup><sub>0.10</sub> |
| Amygdala | Right | 0.04<sup>0.05</sup><sub>0.06</sub> | -0.01<sup>0.01</sup><sub>0.04</sub> |
| Caudate | Left | 0.81<sup>0.81</sup><sub>0.81</sub> | 0.68<sup>0.78</sup><sub>0.84</sub> |
| Caudate | Right | 0.82<sup>0.82</sup><sub>0.82</sub> | 0.52<sup>0.76</sup><sub>0.86</sub> |
| Hippocampus | Left | 0.57<sup>0.58</sup><sub>0.59</sub> | 0.36<sup>0.53</sup><sub>0.64</sub> |
| Hippocampus | Right | 0.58<sup>0.59</sup><sub>0.59</sub> | 0.23<sup>0.50</sup><sub>0.66</sub> |
| Pallidum | Left | 0.55<sup>0.56</sup><sub>0.57</sub> | -0.09<sup>0.30</sup><sub>0.59</sub> |
| Pallidum | Right | 0.54<sup>0.54</sup><sub>0.55</sub> | -0.01<sup>0.38</sup><sub>0.61</sub> |
| Putamen | Left | 0.68<sup>0.69</sup><sub>0.69</sub> | 0.38<sup>0.62</sup><sub>0.75</sub> |
| Putamen | Right | 0.73<sup>0.74</sup><sub>0.74</sub> | 0.41<sup>0.66</sup><sub>0.79</sub> |
| Thalamus | Left | 0.64<sup>0.65</sup><sub>0.65</sub> | -0.08<sup>0.27</sup><sub>0.60</sub> |
| Thalamus | Right | 0.64<sup>0.65</sup><sub>0.65</sub> | -0.09<sup>0.28</sup><sub>0.60</sub> |

#### Appendix A.4.1. Adjusting for ICV

When reporting differences in volume between groups, it is common to adjust for either head size or cerebral volume (Barnes et al., 2010; Mathalon et al., 1993; Voevodskaya et al., 2014). Adjusting by intracranial volume as estimated by FreeSurfer did not improve the intraclass correlations (Table A3).

**Table A3:**
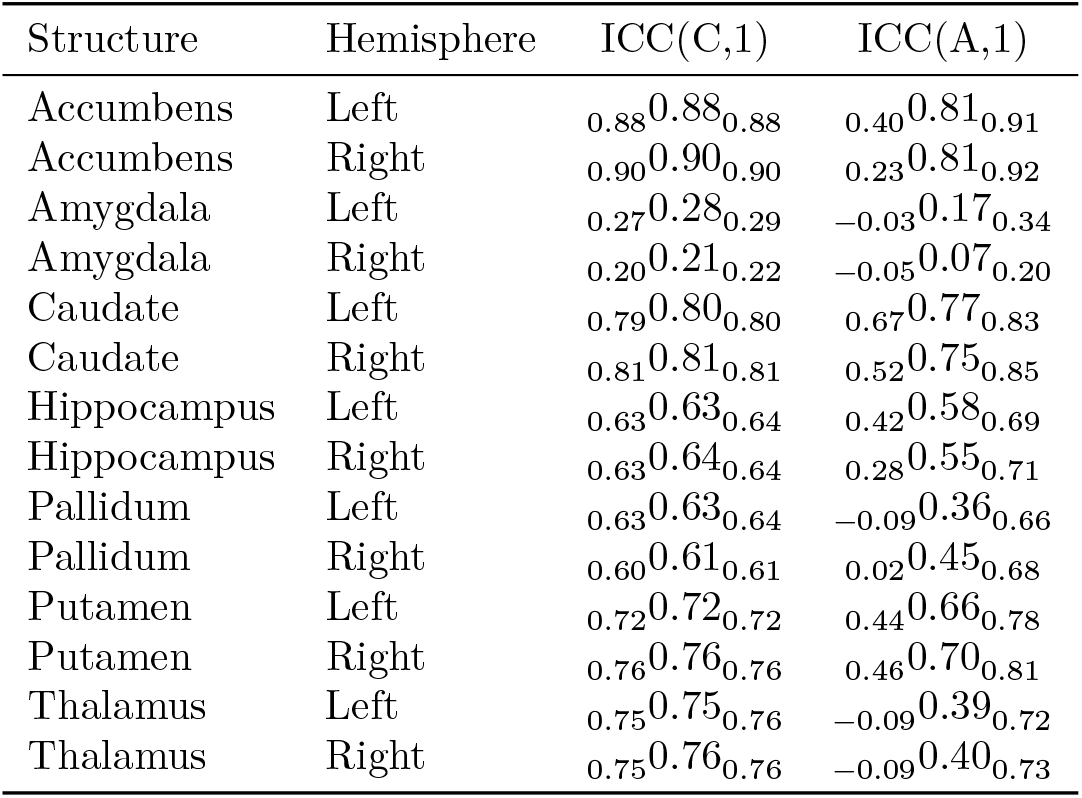
Consistency of Volumes when Adjusting by Intracranial Volume. Adjustments were done by residualizing with respect to ICV. Subscripts indicate 95% confidence intervals.

### Appendix A.5. Simulated Experiments

To simulate hypothetical data, the models described in Section Appendix A.1 and Section Appendix A.7 were used. That is, for a given sample size, 𝐼, true volumes, 𝜆_𝑖_, were sampled for each participant, 𝑖, using a normal distribution, 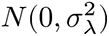. For each participant and tool, 𝑘, mismeasurement of the volumes, were also sampled from a normal distribution, 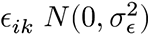. Thus, the measurement for each participant’s volume provided by each tool, denoted 𝑥_𝑖𝑘_, was calculated as the sum of the true volume and the mismeasurement: 𝑥_𝑖𝑘_ = 𝜆_𝑖_ + 𝜖_𝑖𝑘_. Intraclass correlations were calculated across these simulated measurements.

For each simulation, a target variable was generated for each participant, denoted 𝑦_𝑖_𝑘, such that the target variable had a given correlation, 𝜌, with the true volumes in expectation

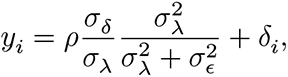

with 𝛿_i_ representing variability in the target variable sampled from a normal distribution, 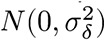.

As described in the main text, simulations considered correlation values equal to 0.01, 0.1, and 0.2. Sample sizes were set to take values between 10 and 100 in steps of 10. For each combination of these parameters, simulated experiments were repeated 1,000,000 times (a high number of repetitions were needed to ensure the stability of estimates of the low rates). Simulated experiments with the UKB data were performed analogously to those with hypothetical data; instead of generating measurements by sampling true volumes and mismeasurement, the measurements sampled from the UKB dataset (UKB samples were chosen with replacement).

In all simulations with artificial data, several parameters described in Section Appendix A. 1 do not influence results after setting the intraclass and interclass correlation and were therefore set to take arbitrary values: 𝜈 = 0, 𝜎_𝛿_ = 1, and 𝜎_𝑐_ = 0. The remaining parameter, 𝜎_𝜆_ was set to 0.019, a value estimated from the full UKB dataset using a linear mixed-effects model and restricted maximum likelihood as implemented by `lme4` (Bates et al., 2015).

To simulate hypothetical data, the models described in the previous sections were used. Sample sizes were set between 10 to 100 in steps of 10. At each combination of parameters, experiments were repeated 1,000,000 times. Simulated experiments with the UKB data were performed analogously to those with hypothetical data (UKB samples were taken with replacement).

Across repetitions, the rates of each effect were estimated by analytic Bayesian methods (binomial likelihood with Beta prior whose shape parameters were set to 1/2). For the effect of “Different Significance”, the posterior was based on the number of simulations in which one correlation exhibited a 𝑝-value less than 0.05 and the other was above 0.05 (successes among relevant simulations) and the number of simulations in which at least one 𝑝-value was below 0.05 (total relevant simulations). The effect of “Different Direction” was calculated similarly, but used simulations in which both correlations had opposite magnitudes out of those in which both were significant. In the main text, ranges of uncertainty refer to 95% equal-tailed intervals and proportions refer to posterior medians.

The 95% equal-tailed interval of the posteriors for the simulations are shown in Figure A3.

### Appendix A.6. Differing Significance Minimally Affected by Increasing Sample Sizes When ICC is Low

As described in the main text, when intraclass correlations were low, increasing the sample size affected the rates of differing significance only minimally Figure 2. This lack of influence can be understood by inspecting the distribution of simulated correlations Figure A4. When the intraclass correlation is low Figure A4, the significance of one correlation is nearly uninformative about the significance of the other (that is, the distributions of the two correlations are nearly circular). Moreover, the power to detect small correlations with even 100 participants is low, and so the power for two tests is very low. But when the intraclass correlation is higher Figure A4, the two correlations cluster, which improves the power of a second test conditioning on one test being significant.

**Figure A4:**
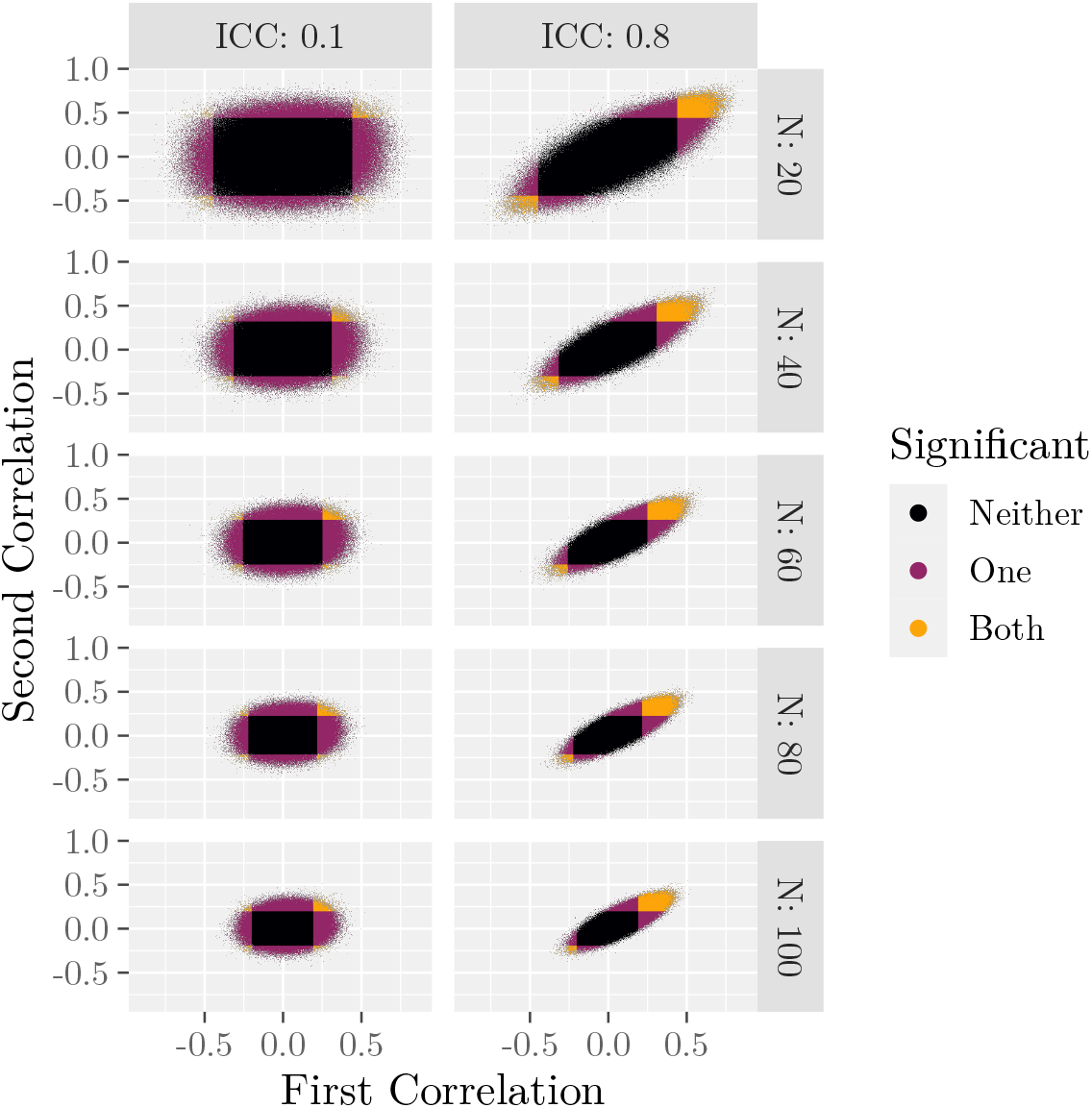
Correlations from Simulations with Artificial Data. Points correspond to simulations and are colored based on statistical significance. The figure shows only a subset of the simulated sample sizes (rows), only the smallest and largest intraclass correlations (columns), and only simulations in which the true effect size (correlation) was 0.1.

### Appendix A.7. Measurement Error and Reduced Correlation Magnitude

Using the model from Appendix A.1, we can build a measurement noise model (e.g., Frost and Thompson, 2000). Call the final target for which we hope to find a relationship with the volume 𝑦. The relationship between that value and the volume was given by an ordinary linear regression with error 𝛿.

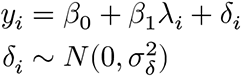

But since we do not know 𝜆, the volumes estimated by the tools are used instead, changing the regression coefficient as follows

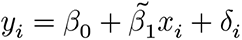

Dilution occurs because the coefficient 𝛽^1^ estimated in this model tends to be closer to zero, decreased by a factor related to the intraclass correlation (Frost and Thompson, 2000).

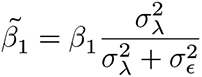

Correspondingly, the desired correlation, 𝜌 = 𝑐𝑜𝑟(𝑦, 𝜆), will also be biased.

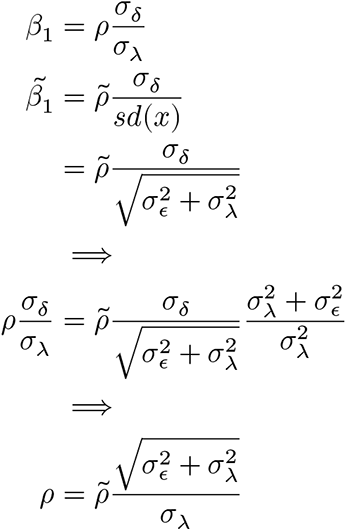

### Appendix A.8. Correlations Between Volumes of the Amygdala and Cognitive Variables

As described in the main text, a set of 50 “cognitive” variables from the UKB was selected for each hemisphere. Selection was based on the rank correlation between the variable and the average (across methods) amygdalar volume. The correlations between those variables and the original volume estimates are shown in Figure A5. For both hemispheres, the magnitude of the correlations with the volumes as reported by FreeSurfer tended to be higher than those reported by FSL.

**Figure A5:**
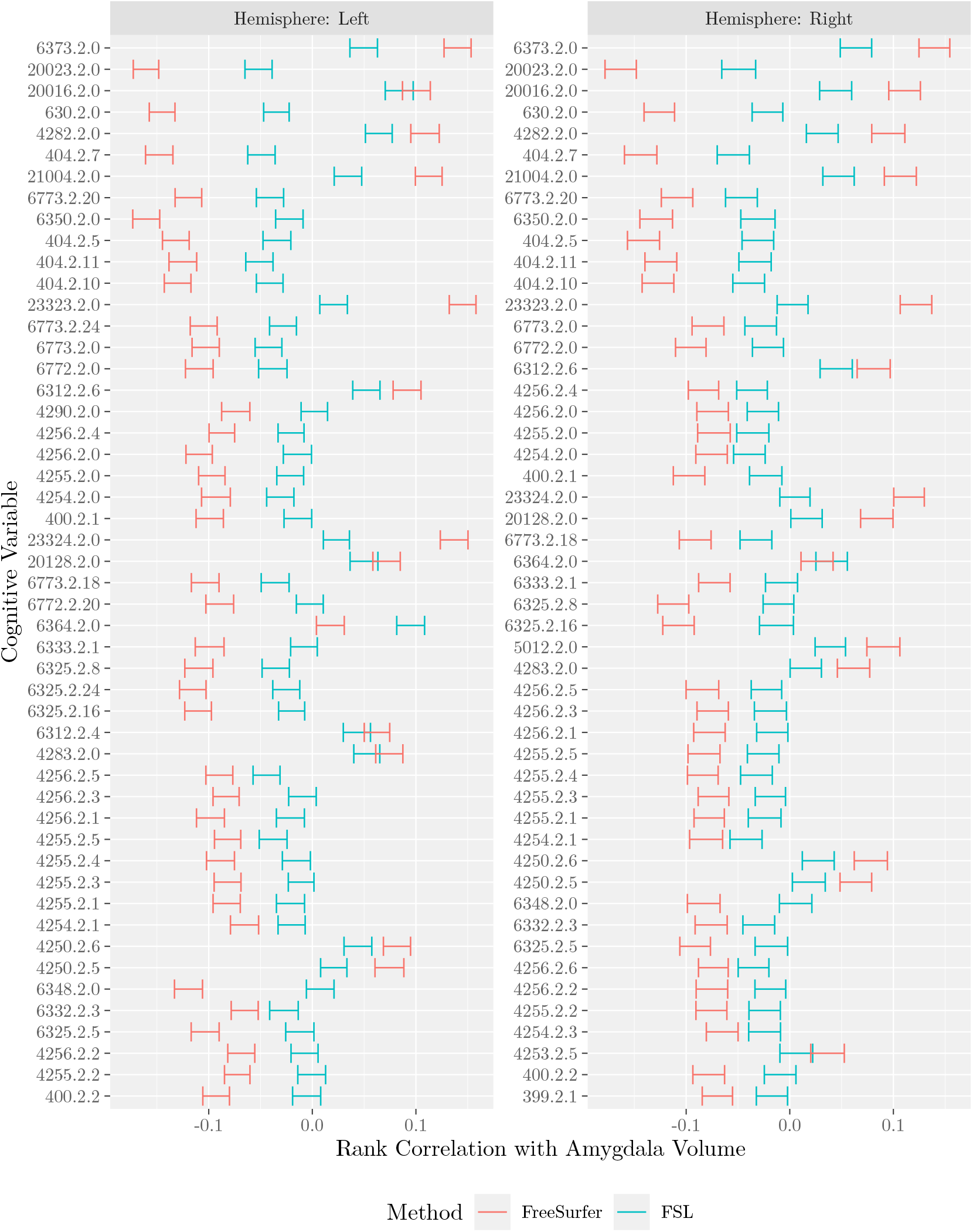
Correlations Between Cognitive Factors and Estimated Volumes of the Left and Right Amygdala. Variables are ordered by decreasing rank correlation using average of left hemisphere volume estimates. Note that variables were selected based on the magnitude of their correlation with amygdalar volumes, which differed across hemispheres, and so the variables in left and right panels differ. Error bars span 95% confidence intervals (bootstrapped with 1000 samples).

